# Meta-analysis refinement of plasma extracellular vesicle composition identifies proplatelet basic protein as a signaling messenger in type 1 diabetes

**DOI:** 10.1101/2022.09.28.509996

**Authors:** Milene C. Vallejo, Soumyadeep Sarkar, Emily C. Elliott, Hayden R. Henry, Fei Huang, Samuel H. Payne, Sasanka Ramanadham, Emily K. Sims, Thomas O. Metz, Raghavendra G. Mirmira, Ernesto S. Nakayasu

## Abstract

Extracellular vesicles (EVs) play important roles in cell-to-cell communication and are potential biomarkers as they carry markers of their derived tissues and disease signatures. However, obtaining pure EV preparations from biofluids is challenging due to contaminants with similar physicochemical properties. Here, we performed a meta-analysis of plasma EV proteomics data deposited in public repositories to refine the protein composition of EVs and investigate potential roles in type 1 diabetes development. With the concept that each purification method yields different proportions of distinct contaminants, we grouped proteins into clusters based on their abundance profiles. This allowed us to separate clusters with classical EV markers, such as CD9, CD40, C63 and CD81, from clusters of well-known contaminants, such as serum albumin, apolipoproteins and components of the complement and coagulation pathways. Two clusters containing a total of 1720 proteins combined were enriched with EV markers and depleted in common contaminants; therefore, they were considered to contain *bona fide* EV components. As possible origins of plasma EVs, these clusters had markers of tissues such as spleen, liver, brain, lungs, pancreas, and blood/immune cells. These clusters were also enriched in cell surface markers CD antigens, and proteins from cell-to-cell communication and signaling pathways, such as chemokine signaling and antigen presentation. We also show that the EV component and type 1 diabetes biomarker, platelet basic protein (PPBP/CXCL7) regulates apoptosis in both beta and macrophage cell lines. Overall, our meta-analysis refined the composition of plasma EVs, reinforcing a primary function as messengers for cell-to-cell communication and signaling. Furthermore, this analysis identifies optimal avenues to target EVs for development of disease biomarkers.

## INTRODUCTION

Extracellular vesicles (EVs) are membrane bilayer-bound particles containing lipids, proteins, nucleic acids and saccharides that are secreted by cells (Raposo & Stoorvogel, 2013). EVs are mainly classified into three groups, exosomes, microvesicles and apoptotic bodies depending on their size and biogenesis. Exosomes range from 30 to 200 nm in diameter and are formed via the endocytic pathway, leading to the formation of multivesicular bodies, which are then fused to the plasma membrane and release EVs. Microvesicles are 100 to 1000 nm in diameter and formed by directly budding from the plasma membrane. Apoptotic bodies are larger EVs (>1000 nm) formed by blebbing of cells undergoing apoptosis (Aguirre *et al*, 2022; Raposo & Stoorvogel, 2013; Salomon *et al*, 2022). EVs have immense potential as biomarkers as they can carry signatures of the tissues of origin and processes affected by disease (Salomon *et al*., 2022). However, a main challenge to study EV function and their potential as disease biomarkers is obtaining pure preparations of EVs from biofluids due to high amounts of contaminants such as lipoproteins and albumin, which have similar physicochemical properties.

Several analytical techniques have been developed for the isolation of EVs, including ultracentrifugation (UC), density gradient ultracentrifugation (DGUC), cushion ultracentrifugation (CUC), polymer-based precipitation (PP), size-exclusion chromatography (SEC) and immunocapture (IC) (Burton *et al*, 2021; Coumans *et al*, 2017). These techniques all have their advantages, but they also suffer from unique contaminant profiles (Burton *et al*., 2021; Coumans *et al*., 2017). For instance, UC co-precipitates particles of similar density while SEC co-fractionates particles of similar sizes (Burton *et al*., 2021; Coumans *et al*., 2017). Sequential purification steps have also been explored (Xu *et al*, 2015). However, they have shown to still be challenging in obtaining pure EV preparations and their labor intensiveness makes them hard to be applied for clinical biomarker studies, which often require large numbers of samples to be properly statistically powered. In an effort to improve the rigor of the EV preparation protocols, the International Society for Extracellular Vesicles (ISEV) developed a guideline with recommendations on experimental design and reporting results (Thery *et al*, 2018). However, there is still a need to systematically evaluate different EV isolation techniques to better understand their performance and the nature of their contaminants.

Our team has been interested on understanding the roles of EVs in type 1 diabetes (T1D) development and their potential as biomarkers of the disease, as we recently discussed in a review article (Aguirre *et al*., 2022). It has been shown that EVs carry molecules, such as microRNAs and chemokines that induces β-cell apoptosis (Javeed *et al*, 2021; Lakhter *et al*, 2018; Zhu *et al*, 2022). However, more systematic studies are needed to refine the composition of EVs and be able to test the role of individual components on disease development.

Here, we performed a meta-analysis of published proteomics data to refine the protein composition of plasma EVs. By systematically analyzing data from multiple studies, metaanalysis enables answering of questions that would unfeasible based on the limited data amounts collected in single projects. We used data from 7 studies of EVs isolated from the blood plasma of healthy human individuals with a combination of various techniques. We reasoned that different purification procedures would result in distinct EV/contaminant ratios that we could use to separate EV-specific proteins from contaminants. Therefore, we clustered proteins based on the abundance profiles, resulting in clusters that were highly enriched with EV markers and others enriched in well-known contaminants. Our results also reveal that plasma EVs are derived from a variety of tissues and that they are enriched for cell signaling proteins. We also showed that the EV component, proplatelet basic protein (PPBP/CXCL7) has a role in cell-to-cell communication in T1D development, regulating apoptosis in both murine beta and macrophage cell lines.

## MATERIALS AND METHODS

### Datasets

Mass spectrometry proteomics datasets of human plasma extracellular vesicles were searched in PRIDE (Perez-Riverol *et al*, 2022) and MassIVE (Choi *et al*, 2020) data repositories of the ProteomeXchange consortium in November and December, 2021, using the key words “extracellular vesicles”, “exosomes”, “microvesicles”, “plasma” and “serum”, following the PRISMA guidelines for meta-analysis (Page *et al*, 2021). Only label-free proteomics data collected on Thermo Orbitrap instruments with data-dependent acquisition were used (**Figure 1**). To reduce potential variability due to pathogenesis processes, only data of extracellular vesicles isolated from plasma of healthy humans were used in our analysis. A list of datasets, their ProteomeXchange identifier, their publication and relevant details of the experimental method are listed in **Table 1**.

**Figure 1.**
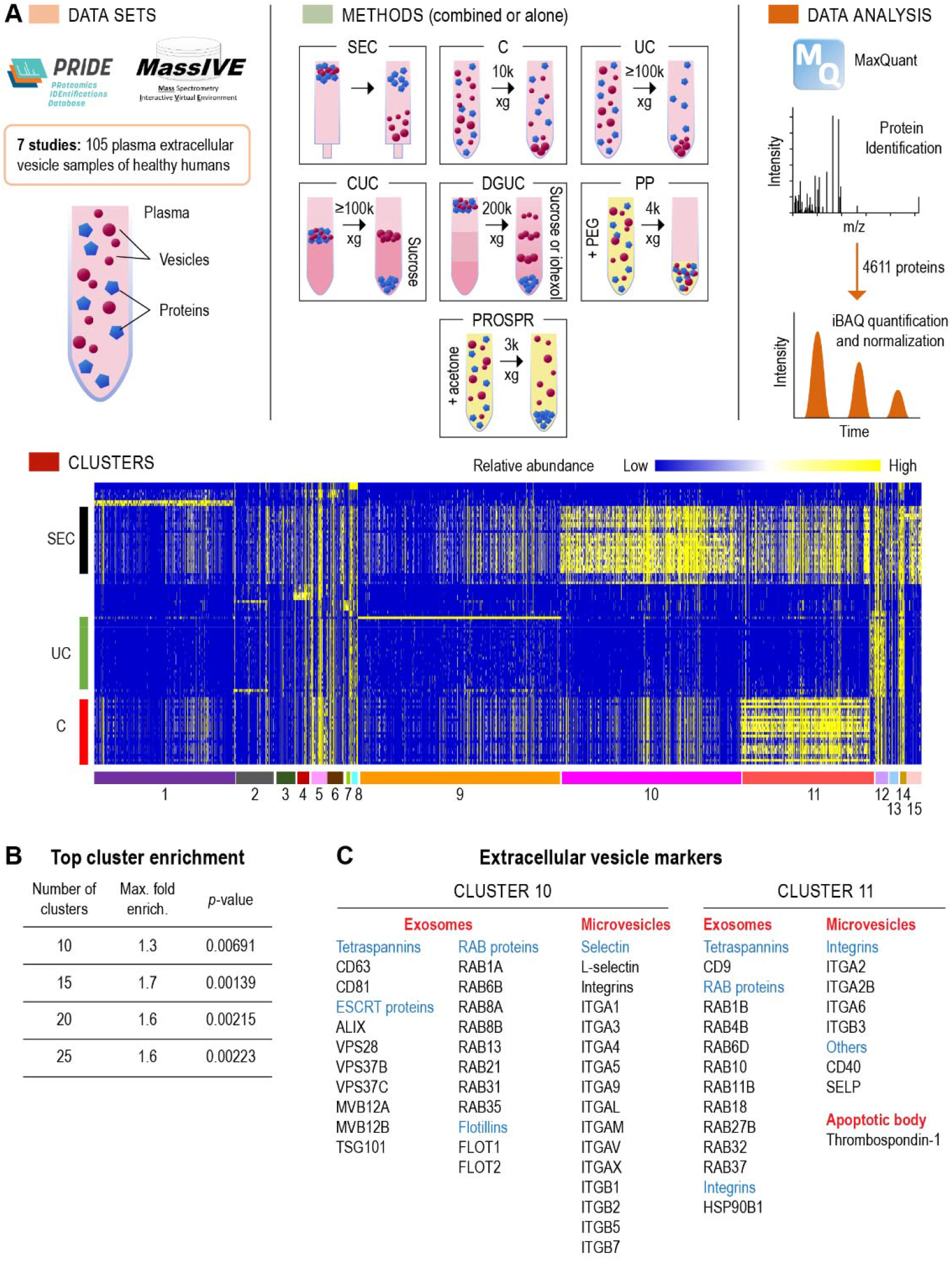
Proteomics meta-analysis of plasma extracellular vesicles (EVs). **(A)** Approach: proteomics data from plasma EV purified with a diversity of methods were downloaded from ProteomeXchange, processed with MaxQuant and submitted to clustering analysis. Abbreviations: C - centrifugation, CUC - cushion ultracentrifugation, DGUC - density gradient ultracentrifugation, DUC - dilution followed by ultracentrifugation, PP - polymer-based precipitation, PROSPR - PRotein Organic Solvent PRecipitation, SEC - size-exclusion chromatography, UC - ultracentrifugation. **(B)** Highest enriched cluster with the top 100 extracellular vesicle proteins from Vesiclepedia when testing different numbers of clusters. (**C**) Classical EV markers found in clusters 10 and 11.

**Table 1.** Characteristics of the proteomics datasets used in the meta-analysis.

| Accession number | Method | Number of samples | Protocol | Ref. |
| --- | --- | --- | --- | --- |
| PXD026483 | Size exclusion chromatography (SEC) | 29 | -Removed cell debris (2,000 xg) and large vesicles (10,000 xg) by centrifugation, for 10 min each<br>-Separated EVs by SEC | (Vander boom *et al*, 2021) |
| PXD024072 | Dilution followed by ultracentrifugation (DUC) | 3 | -Diluted plasma 8x in PBS<br>-Centrifuged at 10,000 xg for 30 min to remove debris<br>-Recovered EVs by centrifugation at 110,000 xg<br>-Wash pellet with PBS/centrifugation at 110,000 xg | (Han *et al*, 2021) |
| PXD016293 | Centrifugation (C) | 25 | -Removed cell debris by centrifuging 2,000 xg for 30 min<br>-Recovered EVs by centrifuging at 10,000 xg for 30 min | (Tunset *et al*, 2020) |
| PXD018301 | Ultracentrifugation (UC) | 32 | -Removed cell debris by centrifugation at 500 x g for 10 min<br>-Removed larger vesicles by centrifugation at 3,000 xg for 20 min, and at 12,000 xg for 20 min<br>-Recovered EVs by centrifugation at 100,000 xg for 70 min<br>-Washed with PBS/centrifugation at 100,000 xg for 70 min | (Hoshino *et al*, 2020) |
| PXD015283 | Polymer-based precipitation (PP)/ Density gradient ultracentrifugation (DGUC) | 3 | -Added equal volume of 20% w/v polyethylene glycol PEG6000<br>-Centrifuged for 15 min at 4,000 xg<br>-Centrifuged on a stepwise iohexol gradient (0-50%) at 200,000 xg for 65 h | (Zhang *et al*, 2020) |
| PXD015283 | PP/SEC | 3 | -Added equal volume of 20% w/v polyethylene glycol PEG6000<br>-Centrifuged for 15 min at 4,000 x g<br>-Separated by SEC | (Zhang *et al.*, 2020) |
| PXD004420 | Cushion ultracentrifugation (CUC_1) | 4 | -Removed cell debris by centrifugation at 300 xg for 10 min<br>-Removed larger vesicles by centrifugation at 10,000 xg for 10 min<br>-Recovered EVs by centrifugation at 100,000 xg for 100 min<br>-Centrifugation of EVs in 40% sucrose cushion at 100,000 xg for 120 min<br>-Wash EV with PBS/centrifugation at 100,000 xg for 120 min | (Haderk *et al*, 2017) |
| PXD002668 | PRotein Organic Solvent PRecipitation (PROSPR) | 3 | -Added 4 volumes of -20°C acetone<br>-Vortexed for a few seconds<br>-Removed vesicle-free proteins by centrifugation at 3,000 xg for 1 min | (Gallart-Palau *et al*, 2015) |
| PXD002668 | Cushion Ultracentrifugation (CUC_2) | 3 | -Removed cell debris by centrifuging 300 xg for 30 min<br>-Recovered EVs by centrifuging at 16,500 xg for 30 min<br>-Centrifuged pellet on a 5.5% sucrose cushion at 200,000 xg for 120 min<br>-Centrifuged the sucrose cushion at 200,000 xg overnight | (Gallart-Palau *et al.*, 2015) |

### Data processing

Data were processed with MaxQuant software (v.1.6.14) (Cox & Mann, 2008) by matching against the human reference proteome database from Uniprot Knowledge Base (downloaded on November 27, 2021). Searching parameters included protein N-terminal acetylation and oxidation of methionine as variable modifications, and carbamidomethylation of cysteine residues as fixed modification when appropriate. Mass shift tolerance was used as the default setting of the software. Only fully tryptic digested peptides were considered, allowing up to two missed cleaved sites per peptide. Each set of data was filtered with 1% false discovery rate (FDR) at both peptide-spectrum and protein levels, resulting in 2% FDR when combined all datasets. Quantification of proteins was done using the intensity-based absolute quantification (iBAQ) method (Schwanhausser *et al*, 2011). The iBAQ values were normalized by the total intensity of the whole sample to calculate the relative copy number of each protein.

### Clustering and enrichment

Clustering analysis of protein abundances was performed with MultiExperiment Viewer - MeV (v 4.9.0) (Chu *et al*, 2008). Missing values were imputed with 1/10 of the smallest value in the whole dataset. Clustering was done by using the K-means clustering method with Pearson correlation and a maximum of 50 iterations. Cluster numbers of 10, 15, 20 and 25 were tested to determine the optimal number. The optimal number was determined by checking the enrichment of the top 100 EV proteins from Vesiclepedia (Pathan *et al*, 2019). Enrichment was calculated based on fold enrichment and statistical significance with Fisher’s exact test.

### Functional-, tissue- and cell-enrichment analysis

Functional-, tissue- and cell-enrichment analyses were done with DAVID (Huang da *et al*, 2009) using the KEGG and Uniprot tissue databases, respectively, using default parameters. Specific tissue markers were mapped by comparing the data against the human tissue proteomics dataset by Jiang et al (Jiang *et al*, 2020). Specific pathways were curated with Vanted (Junker *et al*, 2006).

### Cell growth and apoptosis assay

Murine MIN6 β cell and RAW 264.7 macrophage cell lines were cultivated in DMEM containing 10% FBS and 1% penicillin-streptomycin and maintained at 37 ºC in a 5% CO_2_ atmosphere. Cells were seeded and treated for 24 h with varying concentrations of recombinant platelet basic protein (R&D, cat# 1091-CK-025/CF) followed by a 24 h cytokine cocktail (100 ng/mL IFN-γ: R&D, cat#485-MI-100, 10 ng/mL TNF-α: R&D, Cat#410-MT-010, and 5 ng/mL IL-1β: R&D, cat #401-ML-005) treatment. Apoptosis was measured with the caspase-Glo 3/7 assay (Promega Cat# G8092), following manufacturer recommendations.

## RESULTS

### Meta-analysis and data processing

Plasma EV proteomics datasets were searched in the public repositories associated with the ProteomeXchange consortium, MassIVE and Pride. To keep the datasets consistent, only label-free data collected by data-dependent acquisition in orbitrap mass spectrometers were used in the study. A total of 7 studies were eligible for the meta-analysis. To further make the meta-analysis consistent, we excluded samples associated with disease and only the 105 mass spectrometry datasets from control samples were used (see characteristics in Table with number of samples and purification method) (**Figure 1A**). The data were processed with MaxQuant for peptide/protein identification and quantification, leading to the identification of 4,611 proteins. To compare against different datasets, we used the intensity-based absolute quantification (iBAQ) and normalized by the total intensity of each sample, resulting in the relative protein copies per sample. We performed clustering analysis to group proteins based on their abundance distribution. To have discrete groups, proteins were clustered by the K-means method. The ideal number of clusters were tested in increments of 5 from 10 to 25. We evaluated the ideal number of clusters by performing an enrichment analysis of the top 100 EV proteins from the Vesiclepedia database and considered the ideal number of clusters the one that provides a cluster with the best enrichment with the top 100 EV proteins. A total of 15 clusters showed the best enrichment of the top 100 EV proteins within specific clusters (**Figure 1B, Table 2**). Among the 15 clusters of this analysis, clusters 10 and 11 were significantly enriched with the top 100 EV proteins from Vesiclepedia (1.3 and 1.7 fold, respectively; and p-values of 0.02583 and 0.00139, respectively) and showed a large number of well-known EV markers. Classical exosomal markers, such as tetraspanins CD81 and CD63, were found in cluster 10, whereas CD9 was present in cluster 11. RAB proteins were present in both clusters. Flotillin, ALIX, TSG101 and ESCRT multivesicular-endosome associated proteins, which are frequently identified in exosomes, were segregated into cluster 10. Proteins frequently identified in microvesicles, such as integrins and selectins, were identified in both clusters, while CD40 was detected in cluster 11. Thrombospondin-1, an apoptotic body marker, was associated with cluster 11 (**Figure 1C, Table 2**).

**Table 2.** Enrichment of the top 100 extracellular vesicle proteins from Vesiclepedia across the different protein clusters in the proteomics meta-analysis.

| Cluster | Top 100 | Proteins in cluster | Fold enrichment | p-value | Representative proteins |
| --- | --- | --- | --- | --- | --- |
| 1 | 12 | 777 | 0.6 | 0.01608 | PPIA, NEDD8, ENO1 |
| 2 | 3 | 191 | 0.6 | 0.15103 | H2AC4, CST2, Immunoglobulins |
| 3 | 4 | 144 | 1.1 | 0.1976 | KNG1, VWF, CLEC3B, HBB, FCN3 |
| 4 | 0 | 99 | 0.0 | 0.07808 | APOC1, APOA2, APOA4, APOD, APOA1 |
| 5 | 2 | 90 | 0.9 | 0.27153 | ALB, TF, SERPINA8, F12, F10, F2, A1BG |
| 6 | 3 | 70 | 1.7 | 0.15905 | FGG, F9, C7, C8A, C8B, C9, IGKC |
| 7 | 0 | 52 | 0.0 | 0.26385 | FGB, FBA, CFH |
| 8 | 1 | 49 | 0.8 | 0.36434 | Immunoglobulins |
| 9 | 25 | 1132 | 0.9 | 0.06748 | Ribosome, RNA-binding proteins, tRNA synthases |
| 10 | 32 | 997 | 1.3 | 0.02583 | CD63, CD81, ESCRT proteins, RAB proteins and integrins |
| 11 | 30 | 720 | 1.7 | 0.00139 | CD9, CD40, HLA, RAB proteins, integrins, N-glycosylation proteins |
| 12 | 2 | 92 | 0.9 | 0.26967 | A2M, C3, C5, F13B, C1QA, C1QB, Immunoglobulins |
| 13 | 0 | 72 | 0.0 | 0.1574 | APOB, APOC3, APOC4, APOE, APOF, APOM |
| 14 | 1 | 33 | 1.2 | 0.36889 | HPR, Immunoglobulins |
| 15 | 1 | 93 | 0.4 | 0.22365 | APOC1, APOD, LPA |

### Contaminants

We examined the profile of well-known contaminants (human serum albumin, lipoproteins (based on the apolipoprotein subunits) and proteins from the coagulation and complement pathways) of plasma EV preparations to better understand the performance of different methods. For methods based on centrifugation (centrifugation - C, cushion ultracentrifugation - CUC, dilution followed by ultracentrifugation - DUC and ultracentrifugation - UC) 15% to 35% of the total proteome consisted of proteins from one of the 3 analyzed contaminant classes, most commonly human serum albumin, lipoproteins and coagulation and complement proteins (**Figure 2A**). Compared to the UC, the additional step of CUC failed to reduce the proportion of contaminants in the samples, or to increase the proportion of EV proteins in cluster 10 and 11 (**Figure 2B**). C had the best recovery of proteins from cluster 11 (**Figure 2B**). Protein Organic Solvent PRecipitation (PROSPR) had a similar contamination profile compared to the centrifugation, but lower recovery of clusters 10 and 11. Polyethylene glycol polymer precipitation (PP) followed by density gradient ultracentrifugation (DGUC) highly enriched for lipoproteins, being the apolipoprotein subunits accounting for over 50% of the total protein abundance in the sample (**Figure 2A**). When PP was followed by size-exclusion chromatography (SEC), a similar proportion of contaminants was observed but the serum albumin was the major contaminant (**Figure 2A**). SEC alone had approximately 35% of the sample being among the 3 analyzed contaminants, predominantly lipoproteins (**Figure 2A**). However, SEC alone yielded the best recovery of cluster 10 and the second highest for cluster 11, with these 2 clusters representing approximately 35% of the identified proteome combined (**Figure 2B**).

**Figure 2.**
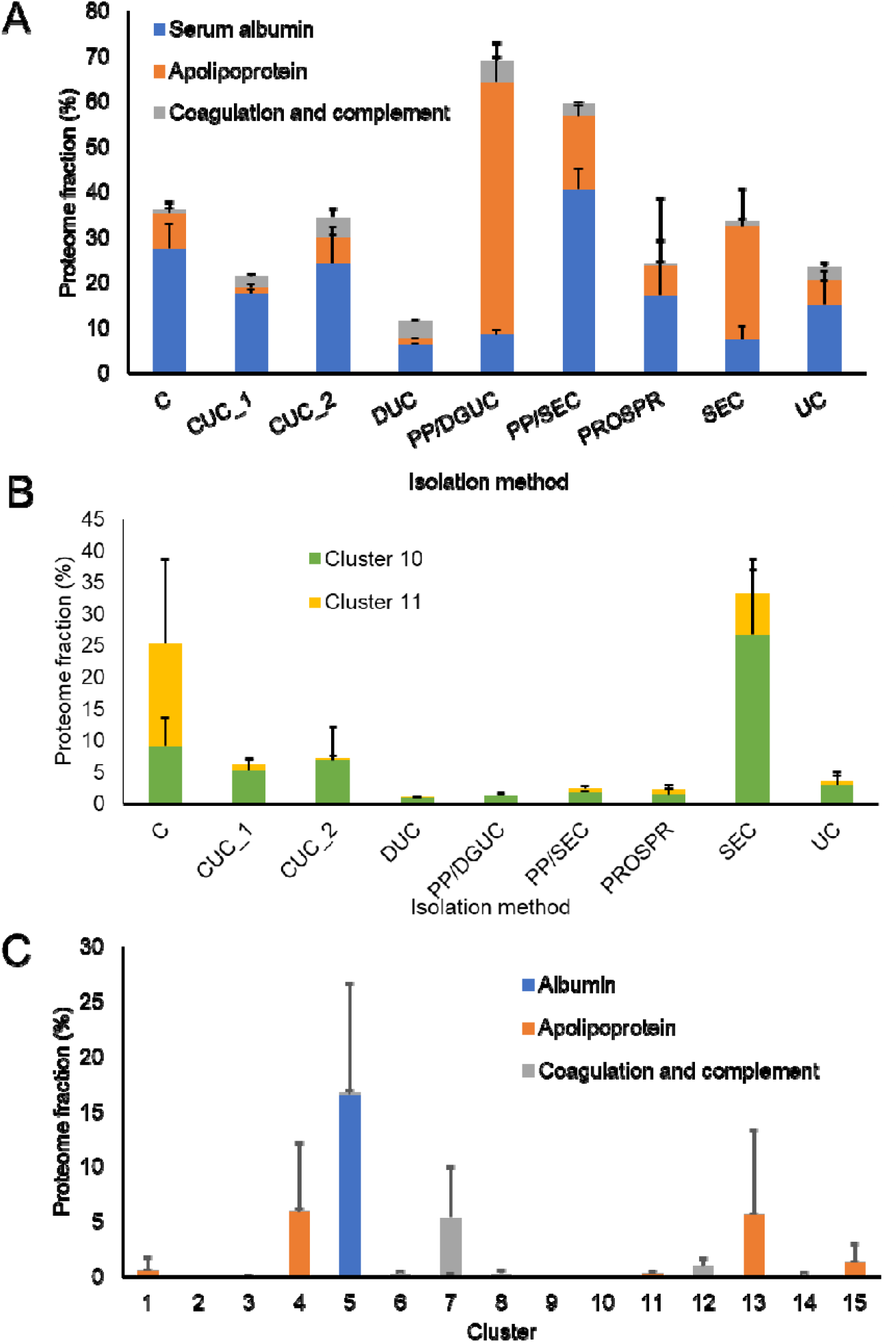
Relative abundance of proteins across different extracellular vesicle purification methods and clusters of the proteome meta-analysis. The proteome samples were quantified by intensity-based absolute quantification (iBAQ) and each protein was normalized by the total sample amount. (**A**) Abundance of common extracellular vesicle preparation contaminants across different purification methods. (**B**) Abundance across different purification methods of clusters 10 and 11, which are enriched in extracellular vesicle markers. (**C**) Abundance of common extracellular vesicle preparation contaminants across different clusters of the proteome meta-analysis. Abbreviations: C - centrifugation, CUC - cushion ultracentrifugation, DGUC - density gradient ultracentrifugation, DUC - dilution followed by ultracentrifugation, PP - polymer-based precipitation, PROSPR - PRotein Organic Solvent PRecipitation, SEC - size-exclusion chromatography, UC - ultracentrifugation.

We examined the profile of serum albumin, lipoproteins and coagulation and complement proteins across 15 clusters. Clusters 10 and 11, which showed the best enrichment in EV proteins, had a low amount of these contaminants in contrast to cluster 5 showed a high amount of albumin, over 15% (**Figure 2C**). Clusters 4, 13 and 15 were abundant in apolipoproteins, ranging from 1.5% to 6.5%, whereas cluster 12 had coagulation complement as the predominant contaminant (0.82%) (**Figure 2C**). These results showed that clusters 10 and 11 were enriched in EV proteins and depleted of common plasma contaminants.

### Origins and markers of plasma EVs

To investigate the tissue and cell origins of the plasma EVs, we performed a tissue- and cell-enrichment analysis using Database for Annotation, Visualization and Integrated Discovery (DAVID). The analysis was based on the Uniprot tissue database and showed that liver, pancreas, kidney, lung, uterus, trachea, heart, colon and skeletal muscle proteins to be enriched in both clusters 10 and 11 (**Figure 3**). Cluster 10 was also enriched in proteins found in tissues from the immune system, spleen, bone marrow, thymus, tonsil, and lymph node, while cluster 11 was enriched in brain (in addition to thalamus, brain cortex, cerebellum), female reproductive tissues (placenta, cervix, uterus, ovary), digestive tissues (tongue, duodenum, colon), skin, kidney, prostate and urinary bladder (**Figure 3**). To find specific tissue markers we compared our data with a comprehensive human tissue proteomics dataset by Snyder et al. In cluster 10, we found specific proteins for tissues that were enriched in the DAVID analysis: spleen (14 proteins), liver (9 proteins), skeletal muscle (5 proteins) and lung (3 proteins). We also found specific proteins from tissues that were not enriched in the DAVID analysis: brain cortex (13 proteins), tibial nerve (5 proteins), small intestine (2 proteins), thyroid (2 proteins), stomach (2 proteins), skin (1 protein), adrenal gland (1 protein) and testis (1 protein) (**Figure 3**). In cluster 11, among the tissue that were enriched in the DAVID analysis we found specific proteins for spleen (11 proteins), liver (4 proteins), skeletal muscle (4 proteins), pancreas (3 proteins) and lung (1 protein). Specific proteins were also found for tissues that were not enriched in the DAVID analysis: small intestine (2 proteins), adrenal gland (1 protein), pituitary (1 protein), minor salivary (1 protein), brain cerebellum (1 protein) and heart atrial (1 protein) (**Figure 3**). These results show that the circulating EVs are derived from a variety of tissues in the body.

**Figure 3.**
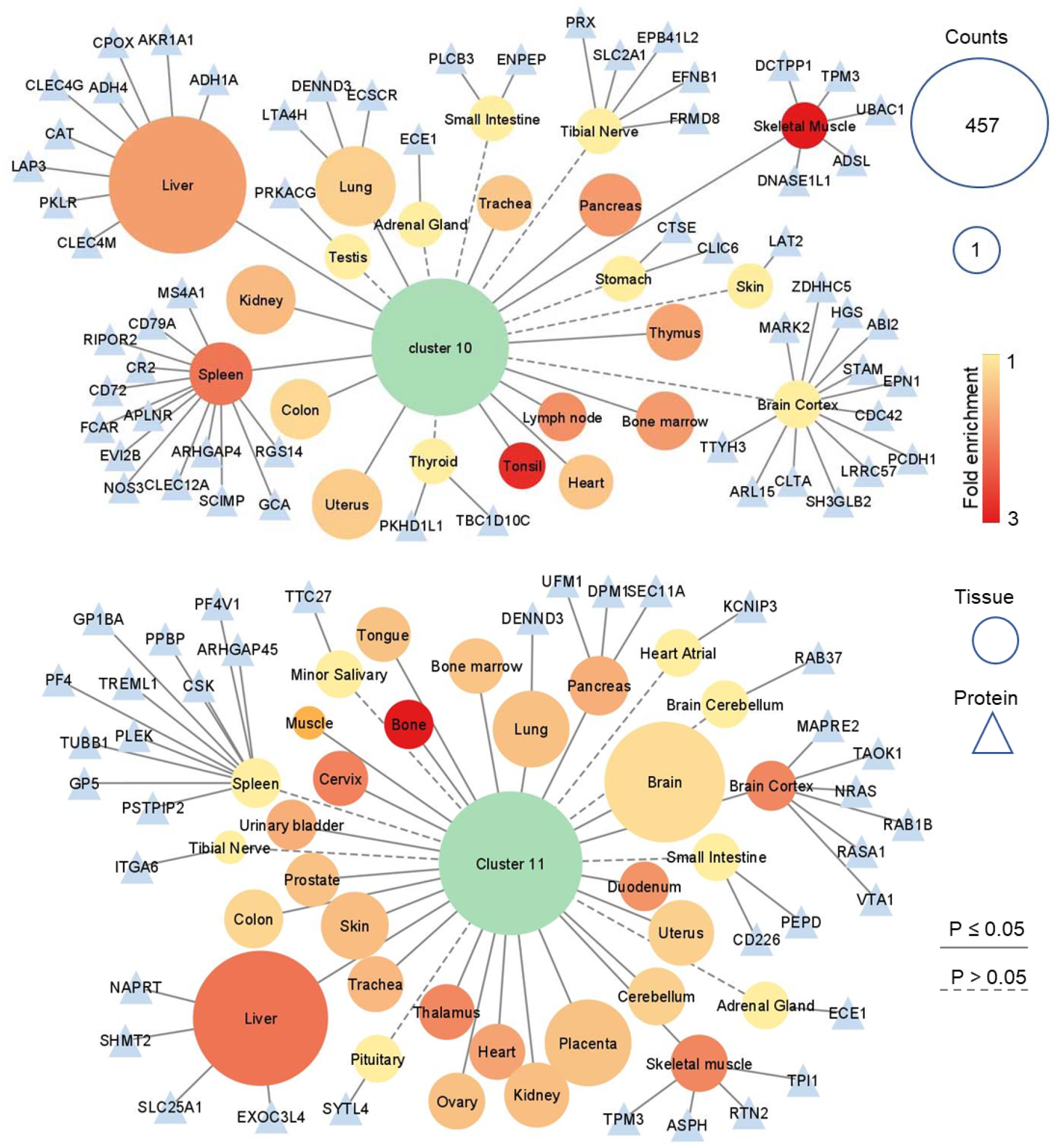
Tissue enrichment and specific tissue markers. Tissues enriched in proteins from clusters 10 and 11 were determined with DAVID. Circle node sizes represents number of proteins and colors, the fold enrichment. Triangles represent tissue-specific proteins determined previously described (Jiang *et al*., 2020), while dotted lines represent tissues with specific markers but not enriched in the DAVID analysis.

We also investigated the possible cellular origins of the plasma EVs using the same approach of tissue-enrichment analysis. Both clusters 10 and 11 were overrepresented in proteins from blood and immune cells, including platelets, erythrocytes, T cells, B cells, monocytes and dendritic cells (**Figure 4A**). The clusters were also enriched in proteins from fibroblasts, keratinocytes, adipocytes and Cajal-Retzius cells, a type of cortical neurons (**Figure 4A**). To look for proteins that can be used as markers of specific cells were looked at the cluster of differentiation (CD) antigens, which are cell surface proteins used distinguish cell populations by immunoassays. We found that CD antigens were enriched in cluster 10, while depleted in other clusters. Out of 188 total identified CD antigens, 133 were present in cluster 10 (**Figure 4B-C**). Cluster 10 had markers of several cells, such as leukocytes (CD45), T cells (CD3), T-helper cells (CD4), cytotoxic T cells (CD8), natural killer cells (CD56), B cells (CD19 and CD20), monocytes/macrophages (CD14 and CD33), granulocytes (CD66b), hematopoietic progenitor cell (CD34), platelets (CD41), red blood cells (CD235a) and endothelial cells (CD146) (**Figure 4C**). These results show that the plasma EV are derived from a variety of cells.

**Figure 4.**
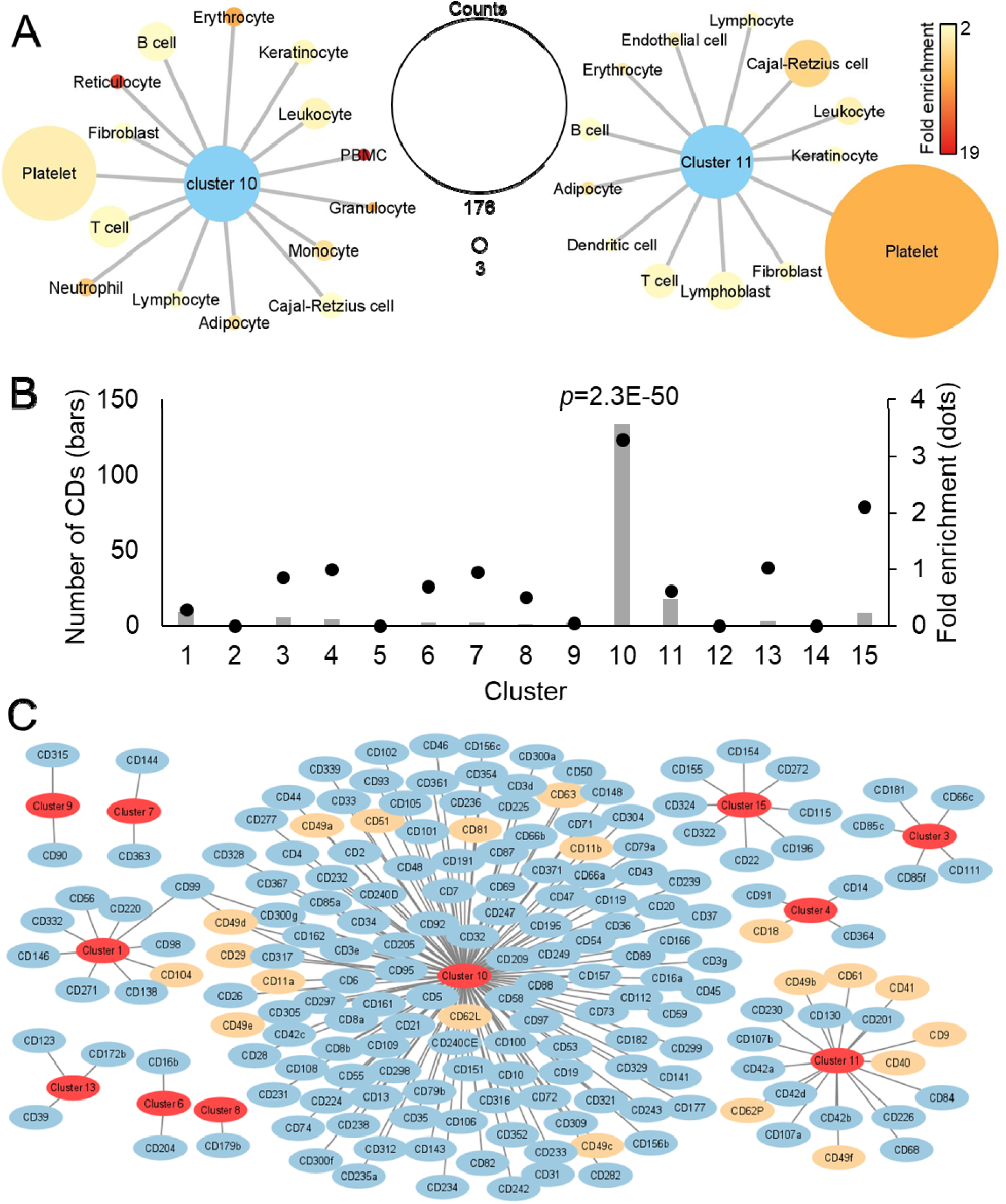
Cellular markers. (A) Cells enriched in proteins from clusters 10 and 11 in the DAVID analysis. Node (circle) sizes represents number of proteins and colors, the fold enrichment. (B) Number of CD antigens and enrichment (compared to the total identified proteins) across different clusters. Significance was determined by Fisher’s exact test. (C) Network of CD antigens across different clusters. Classical extracellular vesicle markers are colored in orange.

### Pathways enriched in EV proteins

To investigate possible functions of plasma EVs, we performed a functional-enrichment analysis using the DAVID tool and the KEGG annotation. Clusters 10 and 11 were enriched in proteins from several cell-to-cell communication pathways, such as antigen processing and presentation, B cell receptor signaling, chemokine signaling, ECM-receptor interaction, T cell receptor signaling and PD-L1 expression and PD-1 checkpoint pathways (**Figure 5**). These clusters were also enriched in proteins from other cell signaling pathways, such as apoptosis, calcium signaling, Fc γ receptor-mediated phagocytosis, insulin secretion, insulin signaling and NOD-like receptor signaling pathways (**Figure 5**). Clusters 10 and 11 were also overrepresented in lipid-mediated signaling pathways, such as estrogen, phosphatidylinositol, phospholipase D and sphingolipid signaling pathways (**Figure 5**). Metabolic proteins, such as from the citrate cycle, cysteine and methionine metabolism, fatty acid metabolism, glycolysis/gluconeogenesis and pentose phosphate pathways, were enriched especially in cluster 11 (**Figure 5**). Proteins from the DNA replication, RNA transcription (RNA polymerase and spliceosome) and protein translation (ribosome and aminoacyl-tRNA biosynthesis) were enriched in cluster 9 (**Figure 5**), which was not enriched with EV markers (**Table 2**). Proteins from the complement and coagulation cascades were highly enriched in clusters 4, 5, 6, 7, 8, 12 and 14, and to a less extent in clusters 3, 10 and 15 (**Figure 5**). These results suggest that EVs may play a role in diverse communication and signaling pathways, in addition to being enriched with specific metabolic pathways. Proteins from DNA replication, RNA transcription, protein translation, and contaminant proteins from complement and coagulation pathways segregated to other clusters not enriched for EV markers, showing that it is possible to distinguish between EV proteins and other contaminants.

**Figure 5.**
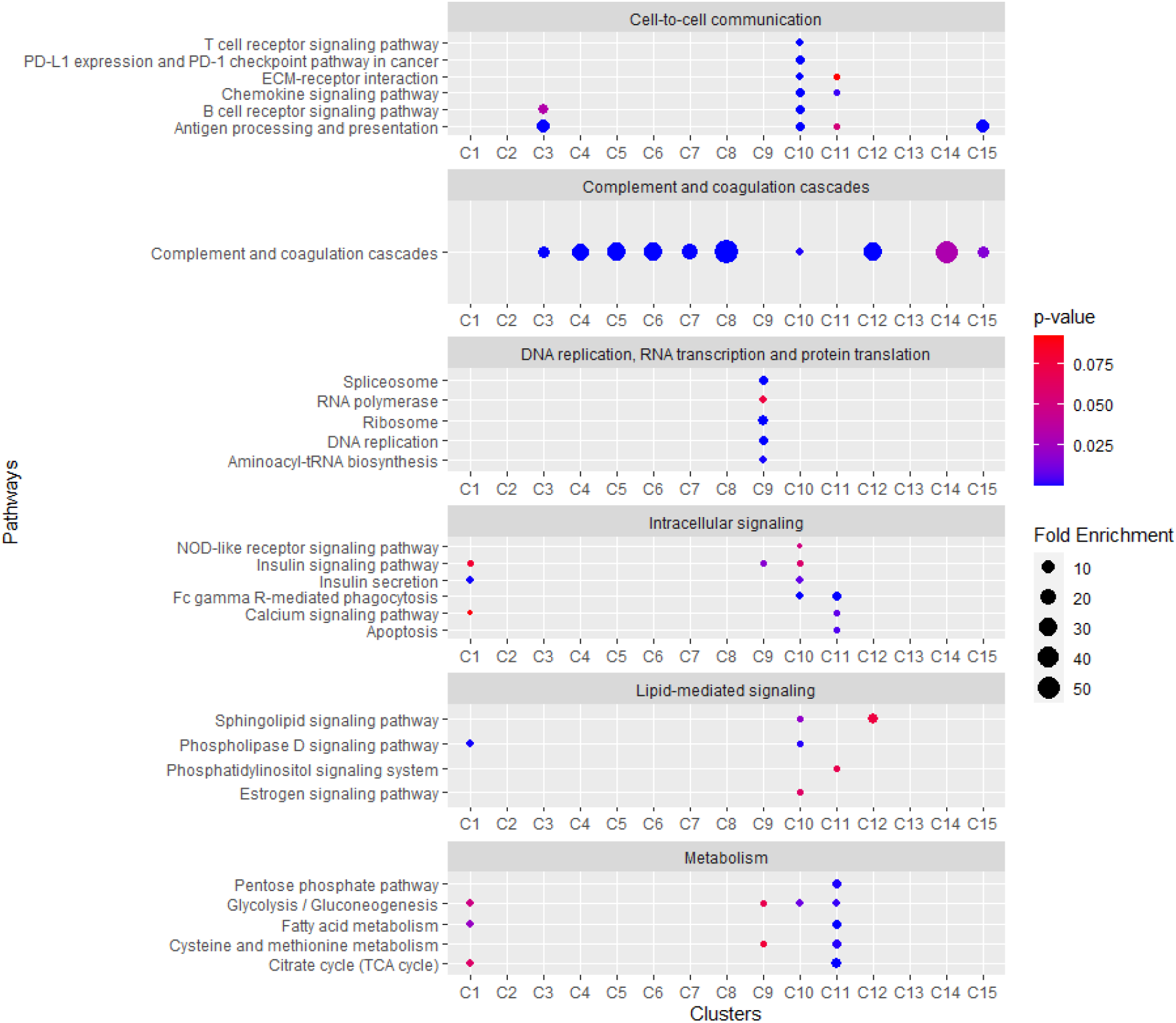
Pathway-enrichment analysis. Distribution of selected pathway enrichment by DAVID analysis across different clusters. The circle sizes represent the fold enrichment and the colors, p-values.

### Chemokine signaling and regulation of apoptosis by the chemokine proplatelet basic protein (PPBP/CXCL7)

Chemokine signaling pathway was highly enriched in proteins from clusters 10 and 11. A total of 7 chemokines and 5 receptors were detected in clusters 10 and 11, along with complete signaling branches, such as the JAK/STAT and SRC/DOCK2/RAC1-2 signaling cascades (**Figure 6A**). We were especially interested in this pathway as many chemokines play a role in T1D development (Eizirik *et al*, 2009). Among the identified chemokines, proplatelet basic protein (PPBP, also known as chemokine CXCL7) (**Figure 6A**) caught our attention since it has been shown to be a T1D biomarker candidate with higher level in plasma prior and at the onset of the disease (Frohnert *et al*, 2020; Zhang *et al*, 2013), but its function in insulitis and β-cell death is unknown. In addition, its main receptor, CXCR2, was also present in the plasma EVs (**Figure 6A**). We investigated if this increase level of plasma PPBP could have a role in regulating cytokine-mediated apoptosis, a key process during T1D development. We performed our experiments in β cells and macrophages because of their role in the disease development. We pre-treated cultured β cells (MIN6) and macrophages (RAW 264.7) with recombinant PPBP and consequently treated with a cytokine cocktail (interferon γ, interleukin β and tumor necrosis factor α) to induce apoptosis. We found that PPBP increase the apoptosis of MIN6 cells, while reduces apoptosis of RAW 264.7 cells (**Figure 6B**). These results show that the EV component PPBP differentially regulate apoptosis in β- and macrophage-cell lines.

**Figure 6.**
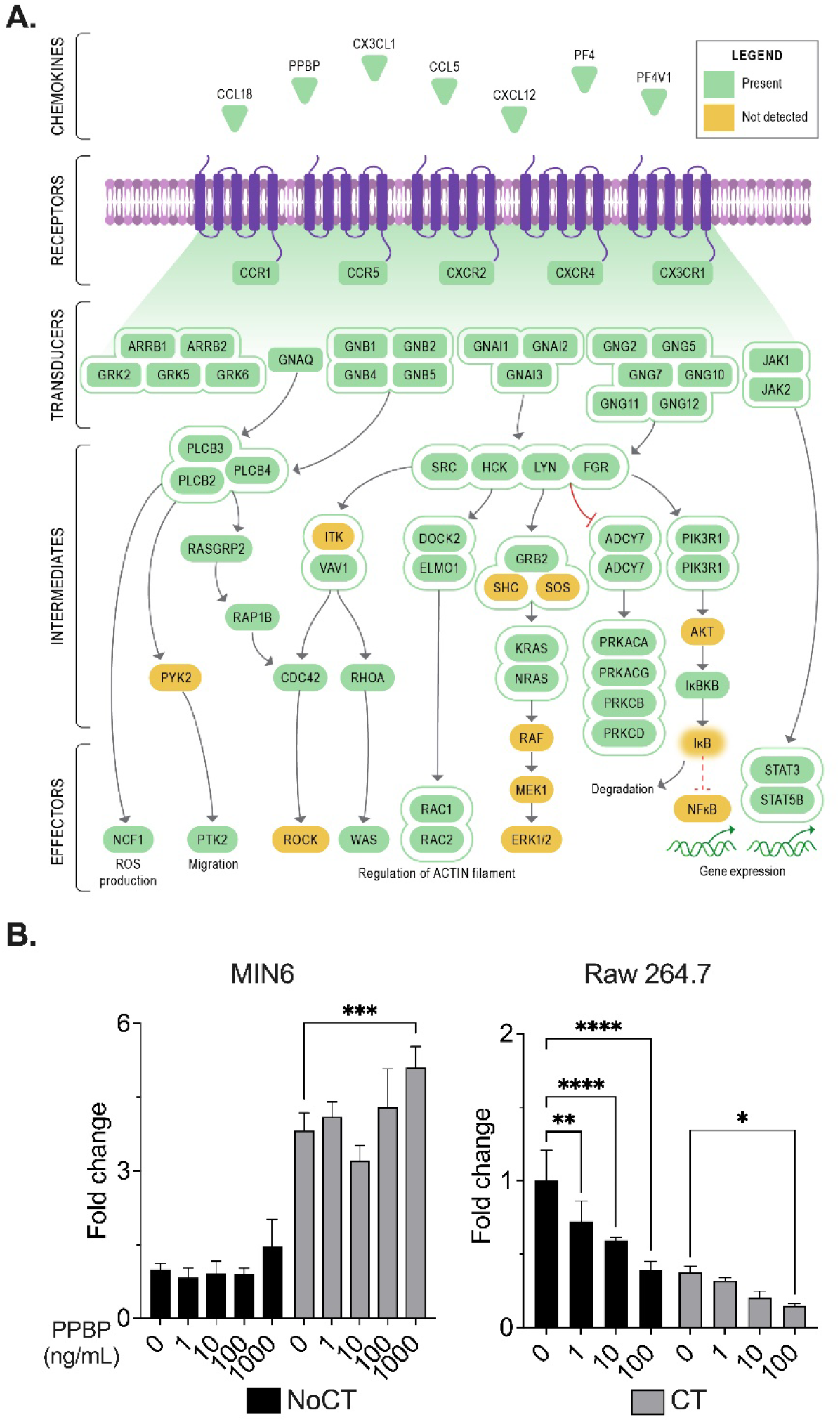
Chemokine signaling components in extracellular vesicles and their activity in cells. (**A**) Chemokine signaling pathway components detected in plasma extracellular vesicles (clusters 10 and 11). (**B**) The Bar graph represents fold change in apoptosis in Raw 264.7 and MIN6 cells pretreated with different concentrations of platelet basic protein (PPBP) for 24 h and then with a cytokine cocktail (100 ng/mL IFN-γ, 10 ng/mL TNF-α, and 5 ng/mL IL-1β). N=4 +SD, Significance by 2-way ANOVA *p<0.05, **p<0.01, ***p<0.001 & ****p<0.0001.

## DISCUSSION

Obtaining highly pure EV preparations is a difficult task. For instance, plasma lipoproteins have similar sizes, and some particles even have similar densities than EVs, making difficult to separate them by size- and density-based techniques (Brennan *et al*, 2020; Burton *et al*., 2021; Coumans *et al*., 2017). Another challenge are highly abundant plasma proteins, such as albumin. Additional purification steps with albumin depletion columns have been explored to remove this contaminant (Palviainen *et al*, 2020). However, multi-step purifications can lead to substantial sample losses, with almost complete EV loss after 3 sequential purification steps (Brennan *et al*., 2020). Therefore, there is a need to better understand the characteristics of different isolation methods towards making better decision on selecting the proper method for each study. By applying a meta-analysis to datasets available from different isolation methods of human plasma, our analysis quantified differences in contaminants across the studied methods and provided insights into strengths and weaknesses of each method.

Centrifugation-based purification methods had similar contamination profiles. Diluting plasma before the centrifugation seems to have a strong effect reducing the contaminants, possibly by diminishing the concentration of proteins and consequently, their aggregation or absorption to EVs. However, the larger volume also affected the recovery of EVs during centrifugation. Addition of purification steps based on the same physical properties (e.g. ultracentrifugation vs. cushion ultracentrifugation) provided no benefit in sample purity. Sequential purification steps have better results when using different physical-chemical properties, such as SEC followed by DGUC, but leads to a low EV recovery (∼1%) (Vergauwen *et al*, 2021). Polymer-based precipitation (PP) had large amounts of lipoprotein contamination, probably due to their mechanism of enriching for EVs by binding to lipids. These findings suggest that understanding of contaminant characteristics is key to understanding limitations of each isolation technique, ways to improve sample preparation quality, and ability to distinguish activities of EVs from those of contaminants. In terms of EV sample recovery, SEC had the best performance recovering proteins from clusters 10 and 11 combined, which is consistent with previous observation that SEC can recover 35% of the plasma EVs (Vergauwen *et al*., 2021). Centrifugation at 10,000 xg showed an enrichment to cluster 11, probably by enriching for EVs with higher density. These observations suggest that for applications that pure materials are not require, such as targeted proteomics analysis, a single step enrichment might be more appropriate than losing most of the material trying to obtain highly purified materials.

In terms of origins of the EVs, we found that they have markers of a variety of tissues and cells. Liver, brain, spleen, lungs, kidneys, pancreas, and bone marrow were among the tissues with the highest enrichment in EV proteins, suggesting that they are main sources of the circulating EVs. This opens a perspective of studying biomarkers for diseases that affect these tissues. Similar to our findings, Muraoka et al showed that plasma EV have proteins that are preferentially expressed in liver, brain, intestine, skeletal muscle and other tissues (Muraoka *et al*, 2022). In terms of cells, EVs were highly enriched in proteins from a variety of cells, especially cells from the immune system, such as lymphocytes, monocytes, neutrophils and dentritic cells. This was accompanied by the presence of cell surface CD antigens, which can be used as cellular or EV markers. Many of these CD antigens have roles in cell signaling and cell-to-cell communication, supporting previous knowledge that EV can participate in those events. A classic example is the antigen presentation and T cell activation by B cell derived EVs (Raposo *et al*, 1996). Our pathway analysis further showed that plasma extracellular vesicles are enriched in cell-to-cell communication and signaling proteins. However, the extent of the enrichment and the completeness of some signaling pathways was unprecedent. This opens the possibility that EVs can not only act as a signaling messenger, but they can also complement cells that lack specific signaling pathways. We also showed that the EV component PPBP to be a regulator of culture β cell and macrophage apoptosis. PPBP has been shown to be a potential T1D biomarker (Frohnert *et al*., 2020; Zhang *et al*., 2013), but its role in the disease development is still poorly understood. PPBP cleavage product, CXCL7, is a major chemokine that primes neutrophil migration by aggregating with platelets and inducing the formation of neutrophil extracellular traps (NETs) (Page & Pitchford, 2013). Neutrophil-platelet aggregates have been found to be abundant in blood of pre-diabetic and recent T1D onset in humans and mice (Popp *et al*, 2022). In addition, NETs were shown to cause β-cell death *in vitro* (Popp *et al*., 2022), further supporting that PPBP might have a role in T1D development.

Overall, we provide an alternative approach for characterizing the EV composition. Our analysis showed that plasma EVs are derived from a variety of tissues and cells and that they are enriched in cell surface markers and cell-to-cell communication molecules. We found that the EV component PPBP controls apoptosis in culture β cells and macrophages and might have a role in T1D development.

## AUTHOR CONTRIBUTIONS

MCV, SHP, SR, EKS, TOM, RGM and ESN designed research study; MVC and ESN performed the meta-analysis and wrote the manuscript; FH, SS, EE, HH performed the experiments and analyzed respective data; SHP, SR, EKS, TOM, RGM and ESN provide guidance and resources to the project. All authors revised the manuscript and gave final approval for publication.

## ACKNOWLEDGEMENTS

This work was supported by the National Institutes of Health, National Institute of Diabetes and Digestive and Kidney Diseases grant U01 DK127786 (to R.G.M, T.O.M. and S.R.), U01 DK127505 (to E.S.N) and R01 DK126444 (to S.R.) S.R. was also supported by UAB-DRC P&F, CDIB Department, and UAB-Comprehensive Diabetes Center. Work was performed in the Environmental Molecular Sciences Laboratory, a U.S. Department of Energy (DOE) national scientific user facility at Pacific Northwest National Laboratory (PNNL) in Richland, WA. Battelle operates PNNL for the U.S. Department of Energy under contract DE-AC05-76RLO01830.

